# Multi-omic Characterization of Pancreatic Cancer-Associated Macrophage Polarization Reveals Deregulated Metabolic Programs Driven by the GMCSF-PI3K Pathway

**DOI:** 10.1101/2021.09.10.459752

**Authors:** Seth M. Boyer, Ho-Joon Lee, Nina G. Steele, Li Zhang, Peter Sajjakulnukit, Anthony Andren, Matthew H. Ward, Venkatesha Basrur, Yaqing Zhang, Alexey I. Nesvizhskii, Marina Pasca di Magliano, Christopher J. Halbrook, Costas A. Lyssiotis

## Abstract

The pancreatic ductal adenocarcinoma (PDA) microenvironment is composed of a variety of cell types and marked by extensive fibrosis and inflammation. Tumor-associated macrophages (TAM) are abundant, and they are important mediators of disease progression and invasion. TAMs are polarized in situ to a tumor promoting and immunosuppressive phenotype via cytokine signaling and metabolic crosstalk from malignant epithelial cells and other components of the tumor microenvironment (TME). However, the specific distinguishing features and functions of TAMs remain poorly defined. Here, we generated tumor-educated macrophages (TEM) *in vitro* and performed detailed, multi-omic characterization (i.e. transcriptomics, proteomics, metabolomics). Our results reveal unique genetic and metabolic signatures of TEMs, the veracity of which were queried against our in-house single cell RNA sequencing (scRNA-seq) dataset of human pancreatic tumors. This analysis identified expression of novel, metabolic TEM markers in human pancreatic TAMs, including ARG1, ACLY, and TXNIP. We then utilized our TEM model system to study the role of mutant Kras signaling in cancer cells on TEM polarization. This revealed an important role for GM-CSF and lactate on TEM polarization, molecules released from cancer cells in a mutant Kras-dependent manner. Lastly, we demonstrate that GM-CSF dysregulates TEM gene expression and metabolism through PI3K-AKT pathway signaling. Collectively, our results define new markers and programs to classify pancreatic TAMs, how these are engaged by cancer cells, and the precise signaling pathways mediating polarization.

## Introduction

Pancreatic cancer is the deadliest major cancer^1^. Early metastasis and insufficient detection methods compound an inability to effectively treat the disease, subjecting patients to a poor prognosis and high mortality rate^2,3^. The tumor microenvironment (TME), composed of a dense fibroinflammatory stroma, has been shown to contribute to the difficulty in treating this disease^4^. In fact, the numbers of malignant cancer cells within pancreatic tumors is typically dwarfed by the immune and fibroblast populations^4^. Accordingly, recent efforts have sought to characterize these non-epithelial components of the TME in pursuit of identifying new and improved detection and treatment modalities. A predominant cell type of interest in the pancreatic TME are tumor-associated macrophages (TAMs), a myeloid cell population that mediates therapeutic resistance and disease aggression^5–11^.

The impact of pancreatic TAMs on tumor growth and aggression has been relatively well established. As the major inflammatory component of solid tumors^12^, TAM abundance correlates with worse response to PDA therapy^5^. The mechanisms by which TAMs mediate this outcome are rather diverse. For example, TAMs promote cancer cell proliferation and metastasis^13^ and protect malignant cells from anti-tumor T-cell activity through immunosuppression^7,11^. TAMs have also been linked to promoting chemoresistance^8^, and recent work by our groups demonstrated that these pancreatic TAMs are capable of directly inhibiting the effect of chemotherapy agent gemcitabine on cancer cells through their release of the pyrimidine nucleoside deoxycytidine^6^. These unique immunosuppressive and metabolic characteristics of TAMs are attributed in part to the phenotypic rewiring macrophages experience in response to the pancreatic TME.

TAMs have long been considered anti-inflammatory “M2-like” macrophages, with *in vitro* models occasionally polarizing naïve macrophages with type-2 cytokines to study TAMs^14^. Although overlap exists between the phenotypes of M2 and tumor-associated macrophages, including oxidative metabolism^6^ and immunosuppressive properties, such as the expression of arginase-1 (Arg1)^15^, the diverse molecular stimuli found throughout the TME polarize TAMs into macrophages with properties not shared with other classical subtypes. The focus of this study is on the mechanistic aspects relating to TAM polarization by directly interrogating direct tumor cell-macrophage interaction.

To recapitulate the signaling and metabolic factors present in the pancreatic TME, we polarized murine bone marrow-derived macrophages (BMDM) *in vitro* with conditioned media from a PDA cell line in which we can regulate the activity of mutant Kras. We refer to macrophages polarized under these conditions as tumor-educated macrophages (TEMs) to distinguish them from TAMs arising in a tumor. We then utilized a systems biology approach integrating our multi-omic profiling (i.e. transcriptomics, proteomics, metabolomics) to define biomarkers for, and the properties of, TEMs. Contrasting this with data from pro-inflammatory “M1-like” and “M2-like” macrophages revealed a panoply of markers and pathways that illustrate the distinct functional characteristics of TEMs relative to classical subtypes. We then queried our in-house, single-cell RNA sequencing (scRNA-seq) dataset^16^ and verified the expression of several of these markers in human pancreatic TAMs, demonstrating persistence of the TAM phenotype across different species and pancreatic cancer models.

Further inquiry into the role of cancer cell mutant Kras activity in TEM polarization led us to observe an important function of a Kras-driven signaling protein (i.e. granulocyte-macrophage colony stimulating factor; GM-CSF) and a metabolite (i.e. lactate) for the expression of several unique TEM markers. Finally, we show that GM-CSF instructs TEM gene expression and metabolism through the PI3K-AKT pathway. Together, these data provide new insights into the crosstalk pathways between cancer cells and macrophages and establish a mechanism by which malignant epithelial cells promote some of the most distinguishing features of TEM function.

### *In vitro* modeling and multi-omic analysis of tumor associated macrophages

To model pancreatic TAMs *in vitro*, we modified the classical BMDM differentiation and polarization paradigm^17^ as follows (**Fig. 1A**). First, we isolated and plated bone marrow in media containing macrophage colony stimulating factor (M-CSF) to differentiate and expand macrophages for 5 days. These naïve macrophages (M0) were then polarized to a tumor-associated phenotype for 2 days in conditioned media from PDA cells. Fresh media was included at a ratio of 1 part to 3 parts conditioned media to account for nutrients consumed by the cancer cells. The resultant *in vitro*-derived cells are herein defined as TEMs, as they do not associate directly with cancer cells. Furthermore, the pancreatic cancer-conditioned media was generated from a cell line (iKras*3) derived from our murine pancreatic tumor model in which reversible mutant Kras expression is under the control of doxycycline (dox)^18^. Growth of these cells in dox drives mutant Kras expression, MAPK pathway activity, and the malignant phenotype *in vitro* and *in vivo*.^18,19^ We also assessed how the removal of Kras from the cancer cells, via dox withdrawal for 3 or 5 days, impacted TEM polarization. In parallel with the TEM polarization strategies, we also polarized M0 macrophages into the canonical *in vitro* phenotypes with 2-day treatment of either lipopolysaccharide (LPS; pro-inflammatory “M1”) or interleukin-4 (IL4; anti-inflammatory “M2”). M0 macrophages were maintained in the naïve state by 2-day treatment with M-CSF (**Fig. 1A**). M1 and M2 phenotypes were independently validated via quantitative polymerase chain reaction (qPCR) of classic pro-inflammatory (Interleukin 12b, *Il12b*; Tumor necrosis factor alpha, *Tnfa*) and anti-inflammatory (Found in inflammatory zone protein 1, *Fizz1*; Chitinase-like 3, *Chil3*; Arginase 1, *Arg1*) genes^20,21^ (**Supplementary Fig. 1A,B**). Importantly, we observed that TEMs did not fit into either the M1 or M2 marker profiles, suggesting that an unbiased approach would be needed to better define PDA-programmed macrophage populations.

**Figure 1.**
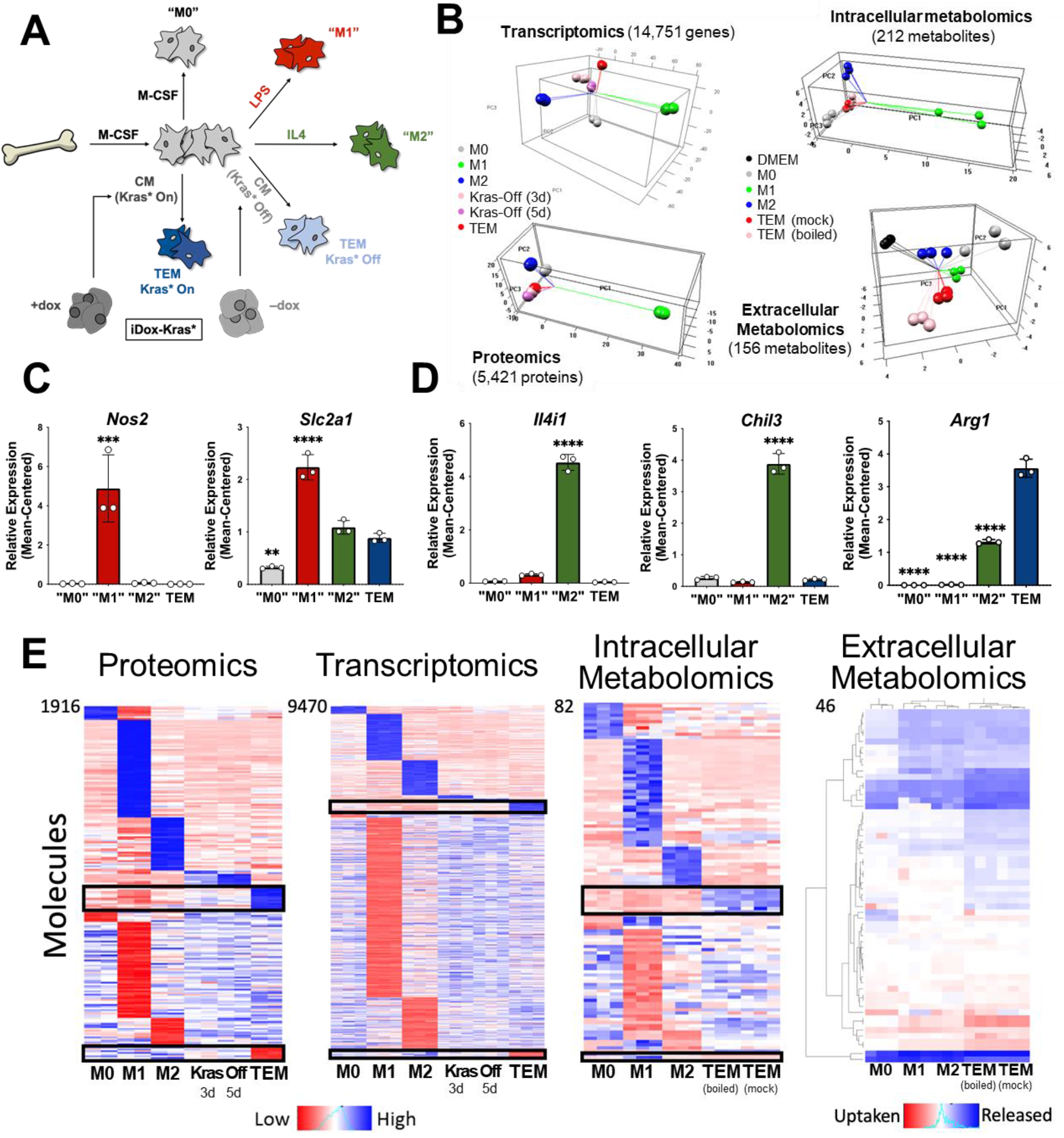
*In vitro* modeling and characterization of pancreatic tumor-educated macrophages. **A**) Schematic of bone marrow-derived macrophage differentiation and polarization. **B**) (Left) Principal component analysis of transcriptomics and proteomics of BMDMs treated with M-CSF (M0), LPS (M1), IL4 (M2), or conditioned media from Kras-Off (3 days or 5 days) or Kras-On (TEM) PDA cells; (Right) intracellular and extracellular metabolomics from media (DMEM), M0, M1, M2 or TEM (Kras-On media was mock-treated or boiled before TEM culture). **C**) RNA-seq-measured mean-centered expression of classical M1 genes *Nos2* and *Slc2a1* across M0, M1, M2, and TEM phenotypes. **D**) RNA-seq-measured mean-centered expression of classical M2 genes *Il4i1, Chil3*, and *Arg1* across M0, M1, M2, and TEM phenotypes. **E**) Heat map array of differential markers of each subtype from proteomics (1,916 proteins), transcriptomics (9,470 transcript), and intracellular (82 metabolites) and extracellular (57 metabolites) metabolomics. Significance comparisons are relative to TEM subtype (* P < 0.05; ** P < 0.01, *** P < 0.001, **** P < 0.0001).

We then performed multi-omic profiling on each of the macrophage subtypes to achieve a comprehensive characterization of genetic and metabolic activity by (i) bulk RNA sequencing (RNA-seq) in triplicates, (ii) proteomic profiling by mass spectrometry in duplicates, and metabolomic analyses on (iii) intracellular and (iv) extracellular metabolites by mass spectrometry in triplicates. Principal component analysis (PCA) from each omics data set demonstrated clustering of the biological replicates reflecting high-quality data (**Fig. 1B**). From this global analysis, we also observed that the M1 subtype has the most distinct molecular profile on all tri-omics levels. The TEM subtype exhibited molecular profiles more similar to the M2 subtype, in line with previous publications from our groups and others^6,15^.

### Metabolism and cytokine signaling are distinctive features of pancreatic TEMs

As a means for further validation, we first directed our attention to known markers of each macrophage subtype in the transcriptomics data. We selected a group of 5 canonical macrophage genes, which were assessed in the primary data. The pro-inflammatory macrophage markers, Nitric Oxide Synthase 2 (*Nos2*) and the glucose transporter *Slc2a1* (GLUT1), displayed increased expression in LPS-treated macrophages compared to the other macrophage subtypes (**Fig. 1C, Supplementary Fig. 1C**). Likewise, IL4-treated macrophages exhibited increased expression of classical anti-inflammatory/tissue remodeling markers, including Interleukin 4 Induced 1 (*Il4i1*), *Arg1*, and *Chil3* (**Fig. 1D, Supplementary Fig. 1B**). Next, we performed differential expression or abundance analysis to identify markers that distinguish each subtype (**Fig. 1E**). The largest number of differential markers occurs in the M1 subtype across all triomics data sets, in agreement with the PCA analysis (**Fig. 1B**).

Given the focus of this study, we narrowed our attention to the TEM subtype. Pathway-centric approaches revealed two prominent features in TEMs relating to (i) cytokine signaling and (ii) metabolism. Differential cytokine signaling is relatively well described for pancreatic TAMs^22,23^. Indeed, C-C Motif Chemokine Receptor 1 (*Ccr1*) and *Ccr5* were among the most highly upregulated genes in TEMs, compared to M0, M1, and M2 macrophages (**Fig. 2A, Supplementary Fig. 1D**). The patterns of differences in mRNA expression were maintained in the proteomics analysis (**Supplementary Fig. 1D**). Of note, our previous assessment of pancreatic TAMs identified CCR1 as a key mediator of immune suppression in pancreatic tumors, corroborating our previous studies.^22^ Further, despite TAMs having long been described as M2-like/anti-inflammatory macrophages, due to their expression of ARG1 and oxidative metabolism^6,15,24^, pancreatic TEMs lack expression of important IL4 targets, demonstrating a clear difference in cellular activity between TEMs and type-2 cytokine-activated macrophages (**Fig. 1D,E, Supplementary Fig. 1B**).

**Figure 2.**
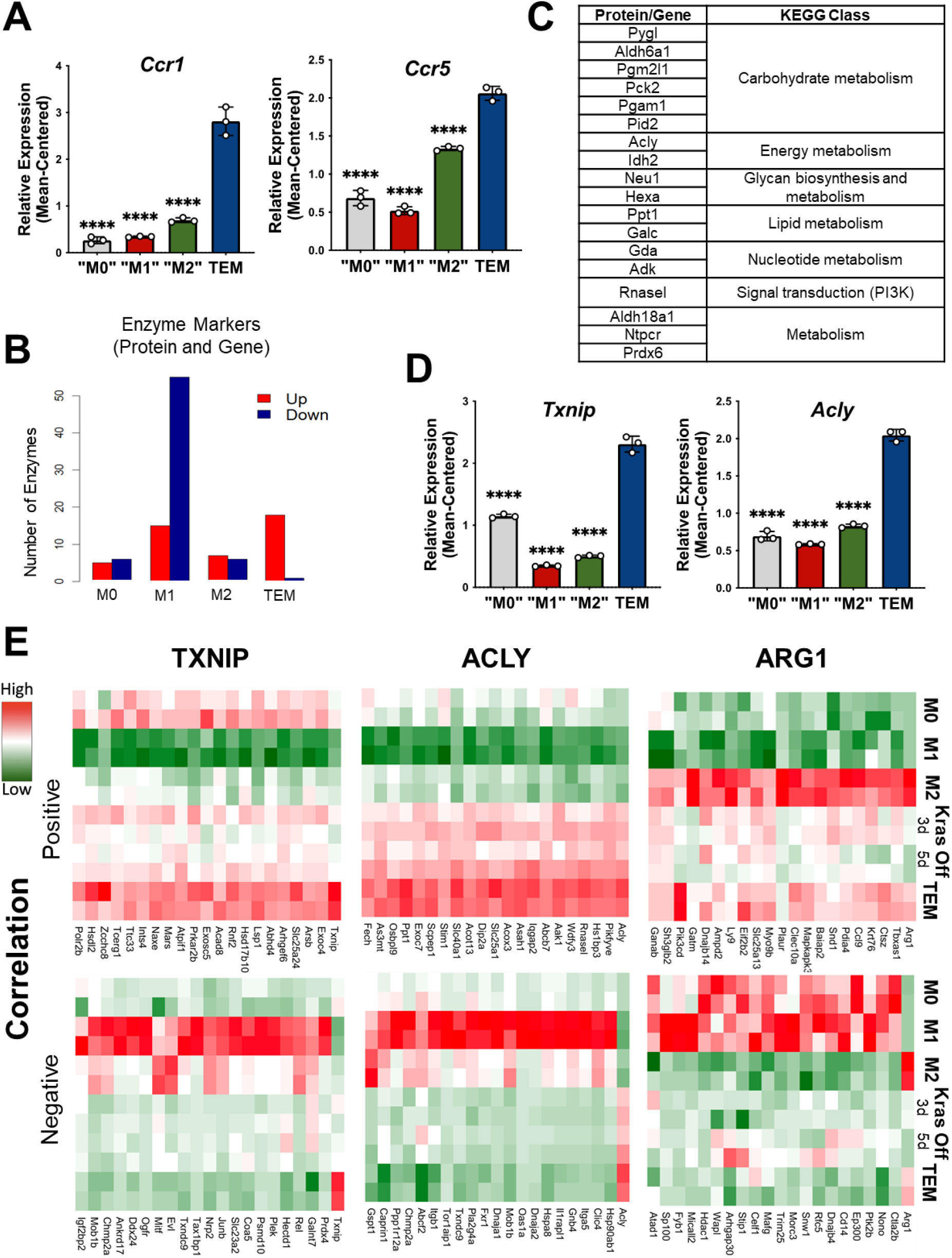
Metabolism and cytokine signaling are distinctive features of pancreatic TEMs. **A**) RNA-seq mean-centered expression of TEM cytokine-related signatures, *Ccr1* and *Ccr5*. **B**) A bar plot of the numbers of up- and down-regulated protein and gene enzyme markers for each subtype. **C**) Table of 18 up-regulated TEM enzyme markers and corresponding KEGG (Kyoto Encyclopedia of Genes and Genomes) class. Note Acly as an enzyme of interest. **D**) RNA-seq-measured mean-centered expression of TEM enzyme signatures *Txnip* and *Acly*. **E**) Heat map of the top-20 positively and negatively correlated proteins for TXNIP, ACLY, and ARG1. Significance comparisons are relative to TEM subtype (* P < 0.05; ** P < 0.01, *** P < 0.001, **** P < 0.0001).

The second differentially enriched pathway in pancreatic TEMs is related to metabolism, and metabolic states have been shown to be key features distinguishing M1 and M2 macrophages.^25^ Indeed, by focusing on enzyme markers from both proteomics and transcriptomics, we find that TEMs have the most up-regulated enzyme markers, while M1 has the largest number of down-regulated enzyme markers (**Figs. 2B,C**).

Among those differential TEM markers, we further narrowed our focus to three markers for follow-up analysis (**Fig. 2D,E, Supplementary Fig. 1B,E**). The first is Thioredoxin-interacting protein (TXNIP), an inhibitor of glucose import^26,27^. Txnip emerged as the top upregulated TEM marker at both the gene and protein levels. The second was ATP Citrate Lyase (ACLY), a well-known enzyme with multi-functional roles in several biological pathways, including serving as a nexus between cellular metabolism and the regulation of gene expression by way of histone acetylation^28^. Finally, we observed *Arg1* to be highly expressed in TEMs (**Fig. 1D**), and even greater than that in “M2-like” macrophages.

Next, we aimed to identify co-expressed proteins or genes of these three markers. Here we focused on correlated proteins, given the more proximal relevance to cellular functions and phenotypes than transcript expression. We selected the top 20 proteins according to both positive and negative correlation with ACLY, ARG1, or TXNIP (**Fig. 2E**). Among those positively correlated with ACLY is SLC25A1, the mitochondrial citrate transporter. Citrate is a substrate of ACLY and highly abundant in TEMs based on our intracellular metabolomics data (**Supplementary Fig. 2A**), suggesting that the pathway of citrate-SLC25A1-ACLY is a TEM signature feature, as has been recently reported in inflammatory macrophages from atherosclerotic plaques^29^. Among the proteins positively correlated with ARG1 is PIK3CD, a component in the PI3K signaling pathway^30^, suggesting its role in TEMs (**Supplementary Fig. 2B**). We also investigated functional associations among those correlated proteins using Search Tool for Retrieval of Interacting Genes/Proteins (STRING)^31^. A particularly strong functional association was found among the TXNIP-correlated proteins, which are mostly involved in metabolism (**Supplementary Fig. 2C**). This is not the case for those *Txnip*-correlated transcripts (**Supplementary Fig. 2D**).

### TEM Markers distinguish pancreatic TAMs in human tumors

To demonstrate biological relevance of the pancreatic TEM phenotype, we queried our in-house single-cell RNA sequencing (scRNA-seq) datasets from human tumors^16^, paying particular attention to the myeloid populations (**Fig. 3A**). We identified expression of several TEM markers in human pancreatic TAMs, such as *Acly* and *Txnip* (**Fig. 3B, Supplementary Fig. 3A**). We are unable to provide sufficient data supporting *Arg1* expression in human pancreatic tumors as it experiences high rates of drop-out during single-cell RNA sequencing. However, we note expression of the strongly *Arg1*-correlated gene, *Pik3cd*, in macrophage populations in human pancreatic tumors (**Fig. 3B, Supplementary Fig. 3A**).

**Figure 3.**
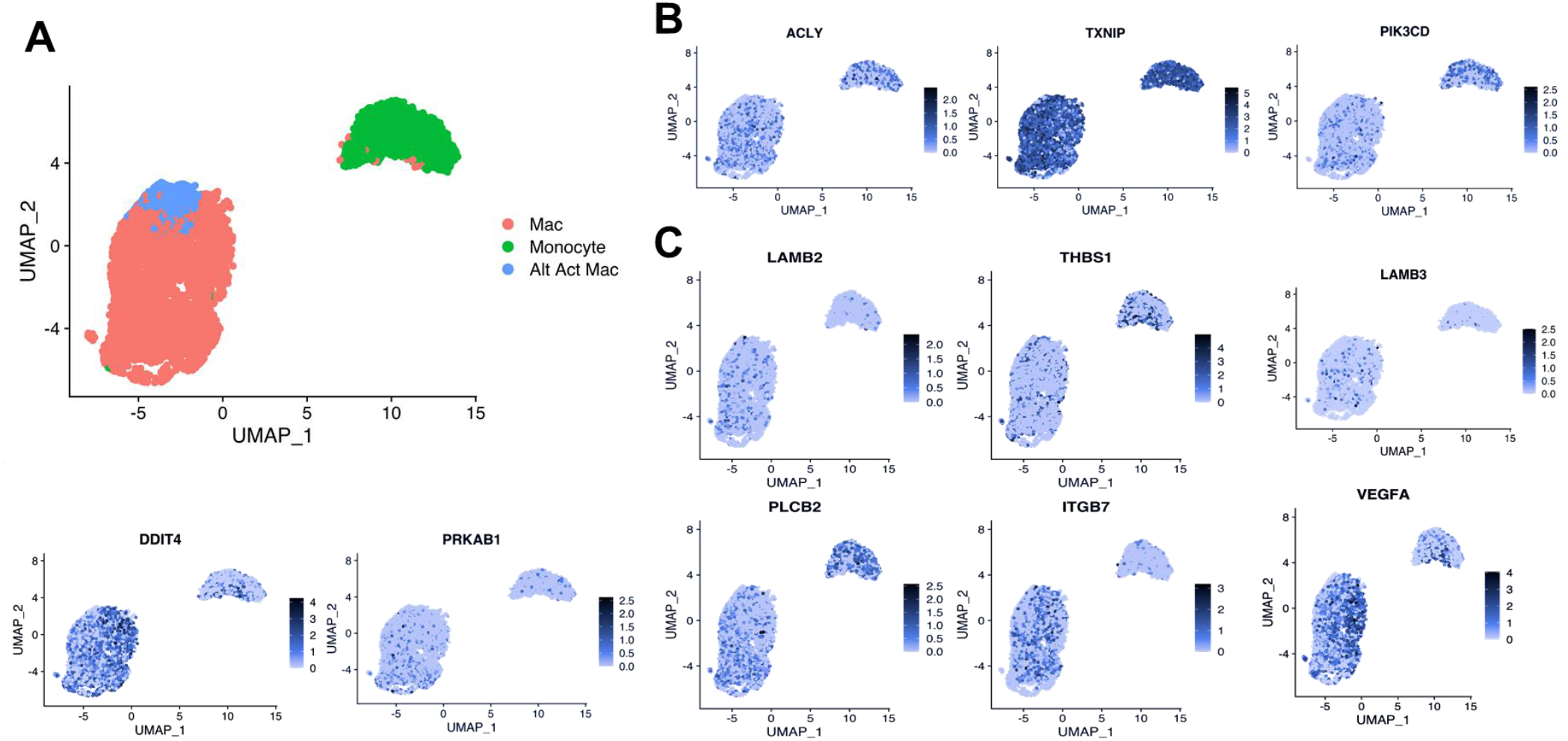
Expression of TEM markers in human pancreatic tumor-associated macrophages. **A**) UMAP plot of myeloid populations in a human pancreatic tumor. **B**) UMAP plots of TEM markers *Acly, Txnip*, and *Pik3cd* in human pancreatic TAMs. **C**) UMAP plots of PI3K-related genes expressed in human pancreatic TAMs (*Ddit4, Prkab1, Lamb3, Lamb2, Thbs1, Vegfa, Plcb2*, and *Itgb7*).

In further support of PI3K relevance in TAMs, we found several PI3K-related TEM signatures (**Supplementary Fig. 2B**) also expressed in human TAMs (**Fig. 3C, Supplementary Fig. 3B**). These data suggest that PI3K signaling in TEMs persists in human TAMs, along with potential contributing factors both upstream and downstream of this signaling pathway.

### Pancreatic TAM polarization is dependent on mutant Kras activity in pancreatic cancer cells

The data from our profiling analyses revealed a clear distinction between the TEMs generated in media from Kras-expressing and Kras-extinguished pancreatic cancer cells (**Fig. 1A,B,E**). As noted in Figure 1, we polarized naïve BMDMs with Kras-On and Kras-Off PDA cell-conditioned media (**Fig. 4A**). Western blot of iKras*3 cell lysates for MAPK pathway activity showed a dramatic decrease in ERK phosphorylation in dox-withheld iKras cells (**Fig. 4B**). We turned our attention to differential markers in Fig. 1E and their expression patterns in Kras-On and 5 day Kras-Off TEMs (**Fig. 4C**). The data revealed broad differences in macrophage gene and protein expression, indicating that inducing mutant Kras in pancreatic cancer cells modifies both the extracellular environment and consequent phenotypes of macrophages exposed to these changes. Specifically, we determined that macrophage expression of *Arg1, Acly, Txnip, Ccr1*, and *Ccr5* were all decreased when PDA cell Kras* was turned off (**Fig. 4D, Supplementary Fig. 4E**).

**Figure 4.**
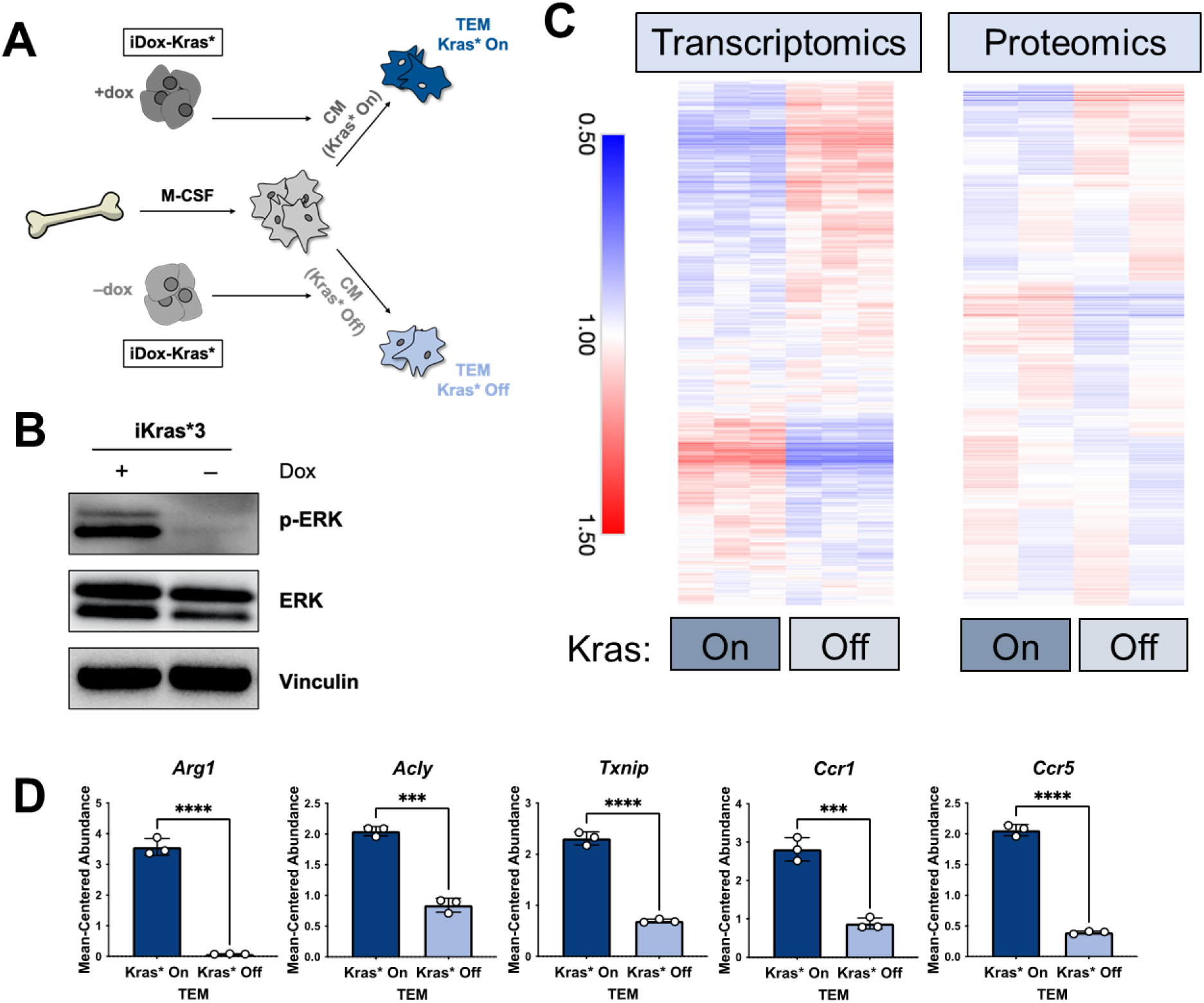
The polarization of pancreatic TEMs is dependent on mutant Kras signaling in pancreatic cancer cells. **A**) Schematic of BMDM differentiation, iKras*3 cell Kras-On and Kras-Off conditioned media generation, and Kras-On and Kras-Off TEM polarization. **B**) Western blot of MAPK pathway proteins ERK and pERK in Kras-expressing and 5 day Kras-extinguished iKras*3 cells. **C**) Transcriptomics and proteomics heat maps of the differential markers in Fig. 1E for Kras-On and 5 day Kras-Off TEMs. **D**) RNA-seq-measured mean-centered expression of TEM signatures Arg1, Acly, Txnip, Ccr1, and Ccr5. (* P < 0.05; ** P < 0.01, *** P < 0.001, **** P < 0.0001).

### Mutant Kras activity in pancreatic cancer cells polarizes TEMs through GM-CSF and lactate

Upon recognizing the importance of PDA mutant Kras signaling for achieving the TEM phenotype, we considered the specific downstream factors of Kras activity that may contribute to TEM polarization. Macrophage expression of Arg1 and Txnip has been shown to be responsive to lactate and extracellular acidification^32,33^. In addition, studies have demonstrated that macrophage expression of Arg1 may be regulated by signaling downstream of granulocyte macrophage colony stimulating factor (GM-CSF)^34^. Furthermore, previous work from our groups and others have also implicated mutant Kras activity in the activation of glycolysis and lactate excretion and the release of GM-CSF^19,35–37^.

Based on these leads, we determined if Kras expression promotes GM-CSF and lactate release in our isogenic, mutant Kras-inducible cell line model and the subsequent role of these factors on TEM polarization. Analysis of GM-CSF expression by qPCR and release by ELISA indicated that loss of mutant Kras expression reduced *Csf2* expression and GM-CSF release by more than 10 and 1000-fold, respectively (**Fig. 5A**). Further, we found that GM-CSF secretion is abundant in two additional murine pancreatic cancer cell lines; i.e. KPC cell lines, KPC7940 and KPCMT3 (**Supplementary Fig. 4A**). Next, we analyzed mutant Kras-dependent extracellular metabolism, including lactate production, by LC/MS-based metabolomics. Metabolome profiling of the spent media from Kras-expressing PDA cells revealed profound alterations to the extracellular metabolome (**Supplementary Fig. 4B,C**), including Kras expression-dependent increase in lactate release (**Fig. 5B**). The Kras-dependent release of lactate was also analyzed and quantitated using an enzymatic assay-based approach (**Supplementary Fig. 4C**).

**Figure 5.**
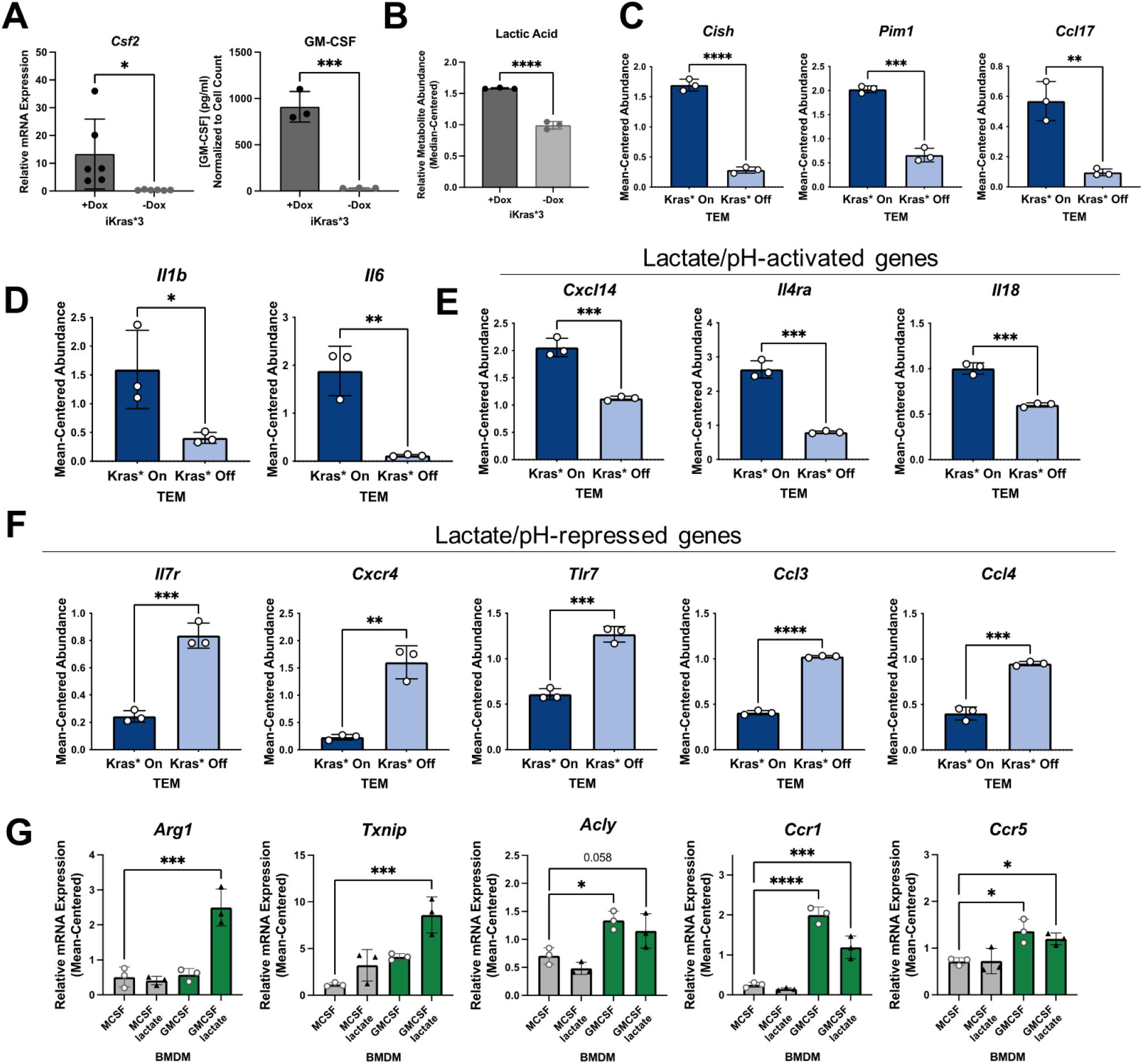
Kras+ PDA polarizes pancreatic TEMs by way of GM-CSF and lactate. **A**) qPCR measurement of *Csf2* expression and ELISA for GM-CSF release. **B**) LC/MS metabolomics-measured median-centered extracellular lactate abundance from Kras-expressing and Kras-extinguished iKras*3 cells. **C**) RNA-seq-measured expression of myeloid GM-CSF-responsive genes, *Cish, Pim1*, and *Ccl17* in Kras-On and Kras-Off TEMs. **D**) RNA-seq-measured expression of lactate-responsive genes, *Il1b* and *Il6* in Kras-On and Kras-Off TEMs. **E**,**F**) RNA-seq-measured expression of genes responsive to acidic extracellular pH in Kras-On and Kras-Off TEMs. **G**) qPCR-measured expression of TEM markers *Arg1, Txnip, Acly, Ccr1*, and *Ccr5* in M0 macrophages treated with either lactate, GM-CSF, or the combination. (* P < 0.05; ** P < 0.01, *** P < 0.001, **** P < 0.0001).

To confirm that these Kras-dependent differences in PDA cell activity impact activation of macrophages, we queried several differentially expressed genes in TEMs that are regulated by PDA mutant Kras and have been documented as responsive to GM-CSF, lactate, or pH. GM-CSF signaling in myeloid cells activates expression of *Cish, Pim1*, and *Ccl17*^38^. Indeed, we demonstrate that expression of these transcripts is significantly upregulated in macrophages exposed to Kras-On PDA cell-conditioned media compared to those exposed to Kras-Off PDA cell-conditioned media (**Fig. 5C**).

*Il1b* and *Il6* were previously identified as lactate-sensitive genes in macrophages^39^. We demonstrate from our RNA-seq data that Kras-On media causes macrophages to express *Il1b* and *Il6* significantly more than macrophages exposed to Kras-Off media. Furthermore, IL6 production has been shown to control macrophage Arg1 expression in an autocrine-paracrine manner^40^, supporting the notion that Kras-dependent increases in lactate production impact the TEM phenotype (**Fig. 5D**).

Lactate is chiefly responsible for acidification of both the tumor microenvironment and the media used in tissue culture; the latter is well appreciated by the yellow shift of the pH sensitive phenol red reagent. Multiple genes expressed by macrophages have been categorized as dependent on extracellular pH, with *Cxcl14, Il4ra*, and *Il18* shown to be increased in acidic extracellular conditions, and *Il7r, Cxcr4, Tlr7, Ccl3*, and *Ccl4* shown to be decreased in acidic conditions^33^. Our RNA-seq data of Kras-On vs. Kras-Off TEMs support these patterns, with each of the aforementioned genes that are increased in acidic conditions upregulated in Kras-On TEMs, and each of these genes that are decreased in acidic conditions downregulated in Kras-On TEMs (**Fig. 5E,F**). These data collectively support the hypothesis that increased cancer cell production of GM-CSF and lactate is dependent on mutant Kras, and that the differential production of mutant Kras-dependent factors modifies both the extracellular environment and subsequent phenotypes of neighboring macrophages.

To further confirm that extracellular GM-CSF and lactate are important contributors to the TEM phenotype, we treated naïve macrophages (M0) for 48 hours with either GM-CSF, lactate, or the combination. A naïve macrophage culture was maintained with M-CSF as a control. In order to more closely mimic the cancer cell-conditioned media, and to assess effects that lactate-induced extracellular acidity may have on TEM polarization, we maintained media supplemented with lactate at a lower pH. We then collected cell lysates and performed qPCR for our TEM markers. These results demonstrated that *Arg1* and *Txnip* are not significantly increased by GM-CSF or lactate as independent treatments. In contrast, these two factors in combination impose a synergistic effect on the expression of both genes (**Fig. 5G**). We also identified increases in *Acly, Ccr1*, and *Ccr5* expression in macrophages treated with GM-CSF. These results, in combination with the increased levels of lactate and GM-CSF observed in Kras-On media, provide strong supporting evidence for the essential role of these factors in TEM polarization.

### PDA-derived GM-CSF promotes TEM polarization through the PI3K-AKT pathway

GM-CSF has pleiotropic effects on signal transduction, dependent on signal strength and context^41,42^. Classic downstream pathways activated by GM-CSF include NFκB, PI3K/AKT and the MAPK pathway. The data presented in Fig. 2E and Supplementary Fig. 2B suggested that TEM polarization was marked by an increase in the PI3K pathway. To test the role of PI3K signaling in TEM polarization downstream of GM-CSF, we activated BMDMs with Kras-On media in the presence or absence of either the pan-PI3K inhibitor, BKM120, the pan-AKT inhibitor, MK-2206, or vehicle control. Western blotting for ARG1 revealed a strong activating role for the PI3K/AKT pathway in pancreatic TEMs (**Fig. 6A**). AKT is known to phosphorylate ACLY, which has been shown to then modify histone acetylation that impacts Arg1 expression in IL4-stimulated macrophages^43^. In support of these findings, we also demonstrate that PI3K/AKT inhibition decreases ACLY phosphorylation in our TEM model (**Fig. 6A, Supplementary Fig. 5A**).

**Figure 6.**
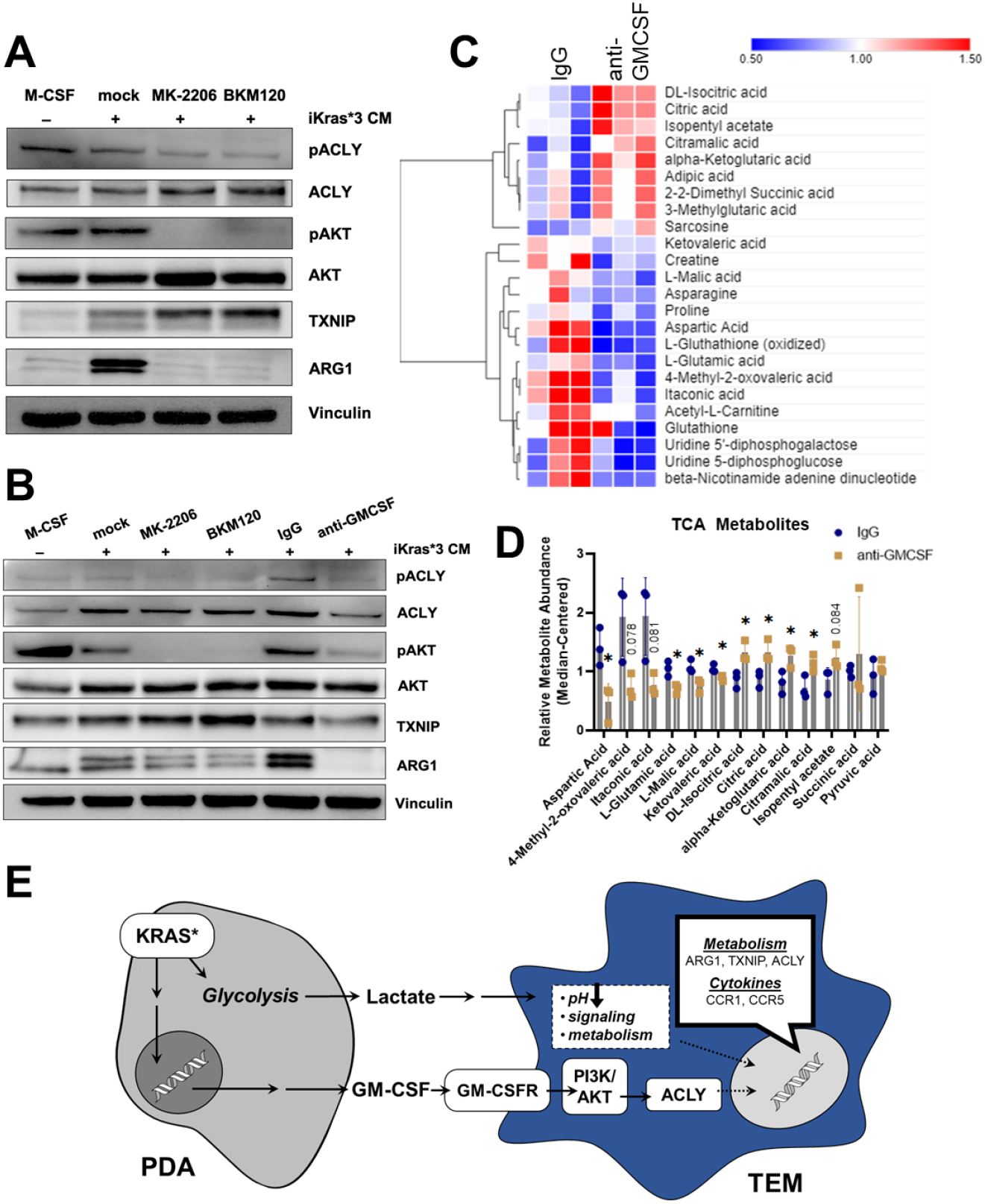
PDA-derived GMCSF promotes TEM polarization and dictates their metabolism through the PI3K-AKT pathway. **A**) Western blot of pACLY/ACLY, pAKT/AKT, TXNIP, and ARG1 in BMDMs treated with either M-CSF, iKras*3 cell-conditioned media + vehicle, iKras*3 cell-conditioned media + MK-2206, or iKras*3 cell-conditioned media + BKM120. **B**) Western blot of pACLY/ACLY, pAKT/AKT, TXNIP, and ARG1 in BMDMs treated with either M-CSF, iKras*3 cell-conditioned media + vehicle, iKras*3 cell-conditioned media + MK-2206, iKras*3 cell-conditioned media + BKM120, iKras*3 cell-conditioned media + IgG control, or iKras*3 cell-conditioned media + GM-CSF-neutralizing antibody. **C**) Heat map of differentially abundant metabolites in TEMs treated with either anti-GMCSF or IgG control. **D**) Bar graph of TCA and related metabolites in TEMs treated with either anti-GMCSF or IgG control. **E**) Schematic of TEM polarization model. (* P < 0.05

Because GM-CSF is secreted by Kras-expressing PDA cells, impacts macrophage Arg1 expression^44^, and activates PI3K^42^, we postulated that cancer cell GM-CSF may be activating macrophage Arg1 expression through the PI3K-AKT pathway. To test this hypothesis, we treated BMDMs with Kras-On conditioned media and either a GM-CSF-neutralizing antibody or IgG control. Indeed, blocking GM-CSF resulted in a dramatic decrease in ARG1 expression, as measured by immunoblotting (**Fig. 6B**). GM-CSF neutralization also resulted in decreased phosphorylation of both AKT and ACLY, confirming that GM-CSF activates macrophage Arg1 expression through PI3K signaling (**Supplementary Fig. 5A**). In addition, blocking GM-CSF led to a modest decrease in expression of ACLY and TXNIP (**Fig. 6B, Supplementary Fig. 5B**), which builds upon our M0 + GM-CSF + lactate qPCR data, in support of an activating role of GM-CSF on enzymatic TEM markers.

Lastly, to analyze how these changes in gene expression impact metabolism, we used our LC/MS-based metabolomics profiling approach in TEMs treated with anti-GM-CSF relative to control antibody (**Fig. 6C**). The anti-GM-CSF--treated groups displayed a trend of decreased sugar phosphorylation, relative to the control conditions (**Supplementary Fig. 5C**). In addition, we observed an increase in citrate, potentially reflecting decreased ACLY activity, in treated cells (**Fig. 6D**). In contrast to citrate, other TCA cycle and associated metabolites, including malate, itaconate, glutamate, and aspartate, were decreased following GM-CSF neutralization (**Fig. 6D**), which suggests that GM-CSF blockade disrupts the TCA cycle and metabolism of associated amino acids. Collectively, these data demonstrate that TEMs are functionally coordinated by GM-CSF stimulation of PI3K signaling in order to maintain their metabolic homeostasis (**Fig. 6E**).

## Discussion

The pancreatic TME consists of a heterogenous mixture of cells and extracellular matrix. TAMs are one of the most abundant cell types in PDA and participate in therapeutic resistance through a variety of mechanisms, including resistance to chemotherapy, immunosuppression, and promotion of tumor growth. However, the factors that contribute to the unique functional properties of TAMs remain insufficiently characterized.

Here, we employed a multi-omic approach to molecularly define pancreatic TAMs. Bulk RNA sequencing, mass spectrometry-based proteomics, and LC/MS-based metabolomics revealed several distinctions between TEMs and classical macrophage subtypes. Our focus on metabolism and cytokine signaling as two primary drivers of cellular function revealed Txnip, Acly, and Arg1 as unique contributors to TEM metabolism. The top 20 proteins correlated with Txnip showed enrichment in metabolism. Acly revealed strong correlation with Slc25a1, the mitochondrial citrate transporter, along with an increase in citrate abundance with respect to other macrophage subtypes, suggesting an important role for this pathway in TEM function. Arg1 was strongly correlated with Pik3cd, a component of the PI3K pathway, suggesting its role in TEM polarization. Indeed, we also observe up-regulation of several PI3K-related genes.

Next, we queried our in-house scRNA-seq dataset of human tumors, from which we observed expression of several important TEM markers in human TAMs. We also note expression of PI3K-related genes in human TAMs, indicating persistence of this pathway in macrophage polarization in clinically relevant models. As confirmation of the general understanding of TAMs, we see that pro-inflammatory markers are not substantially expressed in human TAMs, while anti-inflammatory markers are more abundant.

We then directed our attention to the features of pancreatic cancer cells that drive TEM polarization. Using our isogenic, doxycycline-inducible mutant Kras PDA model, we polarized TEMs with conditioned media from either Kras-expressing or extinguished cells. Indeed, we observed that the most distinct markers of TEM metabolism and cytokine signaling are reliant on Kras expression in PDA cells. Kras is known to impact cancer cell glucose metabolism and growth factor expression. Specifically, lactate and GM-CSF are known to be released from cancer cells in greater abundance when mutant Kras is expressed. By querying our bulk RNA-seq dataset, we observed several GM-CSF- or lactate-responsive genes differentially expressed in Kras-On TEMs compared to Kras-Off TEMs. We then investigated how these factors may affect the expression of significant TEM markers and found that naïve BMDMs treated with GM-CSF displayed increased expression of *Ccr1, Ccr5*, and *Acly*. Further, BMDMs treated with both lactate and GM-CSF displayed increased expression of Arg1 and Txnip. These data suggest that TEM and TAM polarization occurs in response to both metabolic crosstalk and growth factor signaling, and build upon previous reports of the role of tumor-derived lactate in TAM polarization^45^.

In consideration of the GMCSF-PI3K pathway, and the strong correlation between ARG1 and PIK3CD, we treated BMDMs with either a PI3K inhibitor, AKT inhibitor, or GMCSF-neutralizing antibody, and observed that both PI3K-AKT inhibition and GM-CSF neutralization reduced Arg1 expression relative to vehicle and IgG control, respectively. We also note changes in TEM metabolism in response to GM-CSF neutralization, most notably an increase in citrate, which may potentially be correlated with reduced *Acly* expression. Collectively, these data demonstrate an important role for mutant Kras in TEM and TAM polarization, and suggest that mutant Kras exhibits its most significant effects through increased release of GM-CSF and lactate from pancreatic cancer cells. This improved an understanding of epithelial-myeloid communication and distinct features of tumor-associated macrophages will hopefully provide new insights into potential pathways for exploitation to improve pancreatic cancer therapy.

## Materials and Methods

### Cell culture

The doxycycline (dox)-inducible (iKras*3) primary mouse PDA cell line used in this study. was described previously.^7^ Cells were maintained in high-glucose Dulbecco’s modified Eagle medium (DMEM) (Gibco) supplemented with 10% fetal bovine serum (FBS) (Corning) at 37°C. iKras*3 cells were also maintained in 1μg/mL doxycycline. In certain conditions, iKras*3 cells were deprived of doxycycline, for either three or five days before conditioning media, to turn mutant Kras expression off and assess Kras-dependent effects on macrophage polarization.

### Conditioned medium generation

PDA cell-conditioned medium was generated by changing the media of >50% confluent iKras*3 plates, removing media after forty-eight hours, and filtering through a 0.45 μm polyethersulfone membrane (VWR). Fresh media was added at a ratio of 1 part to 3 parts conditioned medium to replenish nutrients consumed by cancer cells. L929 conditioned media was prepared for bone marrow-derived macrophage differentiation, as described.^6^ L929 mouse fibroblasts were maintained in fresh DMEM for 48 hours, after which the conditioned media was filtered through a 0.45 μm polyethersulfone membrane.

### Bone marrow-derived macrophage (BMDM) differentiation

Bone marrow was isolated from the femurs of C57B6/J mice as described^17^ and maintained in macrophage differentiation media (high-glucose DMEM with 10% FBS, penicillin/streptomycin (Gibco), sodium pyruvate (Gibco), and 30% L929 conditioned media) for five days. Media was refreshed on day three, and naïve macrophages were polarized on day five.

### Macrophage polarization

BMDMs were polarized with either 10ng/mL murine macrophage colony-stimulating factor (M-CSF) (Peprotech), 10ng/mL lipopolysaccharide (LPS) (Enzo), 10ng/mL murine interleukin-4 (IL4) (Peprotech), 2ng/mL murine granulocyte-macrophage colony-stimulating factor (GM-CSF), or 75% Kras-On or Kras-Off PDA cell conditioned media. In certain conditions, macrophages were spiked with 5mM lactic acid to assess the effects of extracellular lactate and acidic pH on macrophage gene expression. Each macrophage subtype was polarized from matched biological replicates. Macrophages were maintained in the presence of polarization factors for 48 hours.

### GM-CSF neutralization and PI3K/AKT inhibition

BMDMs were differentiated over 5 days then treated for 48 hours with either 10ng/mL murine M-CSF or 75% Kras-On PDA conditioned media with either vehicle control, 1nM MK-2206 (Selleck Chemicals), 1nM BKM120 (Selleck Chemicals), 1ug/ml anti-GM-CSF neutralizing antibody (BioLegend), or IgG control. Compounds were maintained in dimethyl sulfoxide (DMSO). Macrophages polarized in the presence of the PI3K and AKT inhibitors were pretreated with the respective compound for thirty minutes.

### RNA isolation and reverse transcription

Polarized BMDMs were lysed with RLT Plus buffer with β-mercaptoethanol, lysates were homogenized using a Qiashredder, and RNA samples were isolated according to the RNeasy Plus Mini Kit (Qiagen) protocol, which included gDNA eliminator spin columns. All RNA samples were tested for concentration and purity via NanoDrop (Thermo Scientific). RNA samples were stored in −80°C until needed for reverse transcription. Complementary DNA (cDNA) reverse transcription was performed following the iScript cDNA Synthesis kit protocol (BioRad), and cDNA samples were used for quantitative polymerase chain reaction (qPCR).

### Western Blotting

Cells were lysed in radioimmunoprecipitation assay (RIPA) buffer (Sigma Aldrich) and supplemented with phosphatase inhibitor (Sigma Aldrich) and EDTA-free protease inhibitor (Sigma Aldrich). Lysates were quantified by BCA assay (Thermo Fisher Scientific Inc.), and equivalent protein amounts were run onto SDS-PAGE gels. Proteins were transferred from the SDS-PAGE gel to an Immobilon-FL PVDF membrane, blocked, and incubated with primary antibodies. After washing, membranes were incubated in secondary antibody, washed, then exposed on a Biorad Chemidoc with West Pico (Thermo Fisher Scientific) or West Femto ECL (Thermo Fisher Scientific). Quantitation was performed using Image Lab software.

| Protein | Antibody Name | Catalog # | Company | Dilution |
| --- | --- | --- | --- | --- |
| ACLY | ATP-Citrate Lyase Antibody | #4332 | Cell Signaling | *1:1000 |
| p-ACLY | Phospho-ATP-Citrate Lyase (Ser455) Antibody | #4331 | Cell Signaling | *1:500 |
| AKT | Akt Antibody | #9272 | Cell Signaling | *1:1000 |
| p-AKT | Phospho-Akt (Ser473) (D9E) XP® Rabbit mAb | #4060 | Cell Signaling | *1:1000 |
| ARG1 | Arginase-1 (D4E3M™) XP® Rabbit mAb | #93668 | Cell Signaling | *1:1000 |
| TXNIP | TXNIP (D5F3E) Rabbit mAb | #14715 | Cell Signaling | *1:1000 |
| pERK | Phospho-p44/42 MAPK (Erk1/2) (Thr202/Tyr204) (E10) Mouse mAb | #9106 | Cell Signaling | *1:1000 |
| ERK | p44/42 MAPK (Erk1/2) (137F5) Rabbit mAb | #4695 | Cell Signaling | *1:1000 |
| Vinculin | Vinculin (E1E9V) XP | #13901 | Cell Signaling | *1:5000 |
| Anti-rabbit IgG HRP-linked Secondary | Anti-rabbit IgG, HRP-linked Antibody | #7074 | Cell Signaling | *1:5000 |
| Anti-mouse IgG HRP-linked Secondary | Anti-mouse IgG, HRP-linked Antibody | #7076 | Cell Signaling | *1:5000 |

### Lactate measurement

Lactate measurements were carried out using the lactate fluorescence assay kit (Biovision #K607). Assays were performed according to the manufacturer’s instructions. Lactate levels were measured using a SpectraMax M3 Microplate reader (Molecular Devices).

### RNA sequencing (RNA-Seq) and data analysis

RNA sequencing and data analysis were performed as described.^22^. Upon isolation of RNA samples, and determination of RNA concentration and quality, the University of Michigan Sequencing Core prepared strand mRNA libraries that were sequenced using 50-cycle paired-end reads via a HiSeq 4000 (Illumina) sequencing system. Raw data were generated and analyzed by the University of Michigan Bioinformatics Core. A quality control (QC) was performed using FastQC software (Babraham Bioinformatics) for both pre- and post-alignment. Raw sequencing reads were aligned to the University of California Santa Cruz (UCSC) mm10 assembly mouse genome browser with Bowtie2 and TopHat tools of the Tuxedo suite RNA-Seq alignment software. Quantification of gene expression and differential expression analysis were performed with HTSeq and DESeq2 software systems, respectively. Differentially expressed genes were defined by a false discovery rate of 0.05 and a fold change of expression ≥1.5. Relative expression was graphed as mean-centered abundance, in which each sample’s raw expression value was divided by the mean expression value of all samples.

### Quantitative polymerase chain reaction (qPCR)

Samples for quantitative polymerase chain reaction (qPCR) were prepared with 1x Fast SYBR Green PCR master mix (Applied Biosystems). Primers were optimized for amplification under the following conditions: 95°C for ten minutes, followed by forty cycles of 95°C for fifteen seconds and 60°C for one minute. Melt curve analysis was performed for all samples upon completion of amplification. Hypoxanthine phosphoribosyltransferase (*Hprt1*) primer was used as a reference gene. Relative quantification was calculated using the 2^-ΔΔCT^ method, in which the cycle threshold (CT) value of a target sample’s target gene is normalized to the expression of a reference gene in both a reference sample and the target sample.

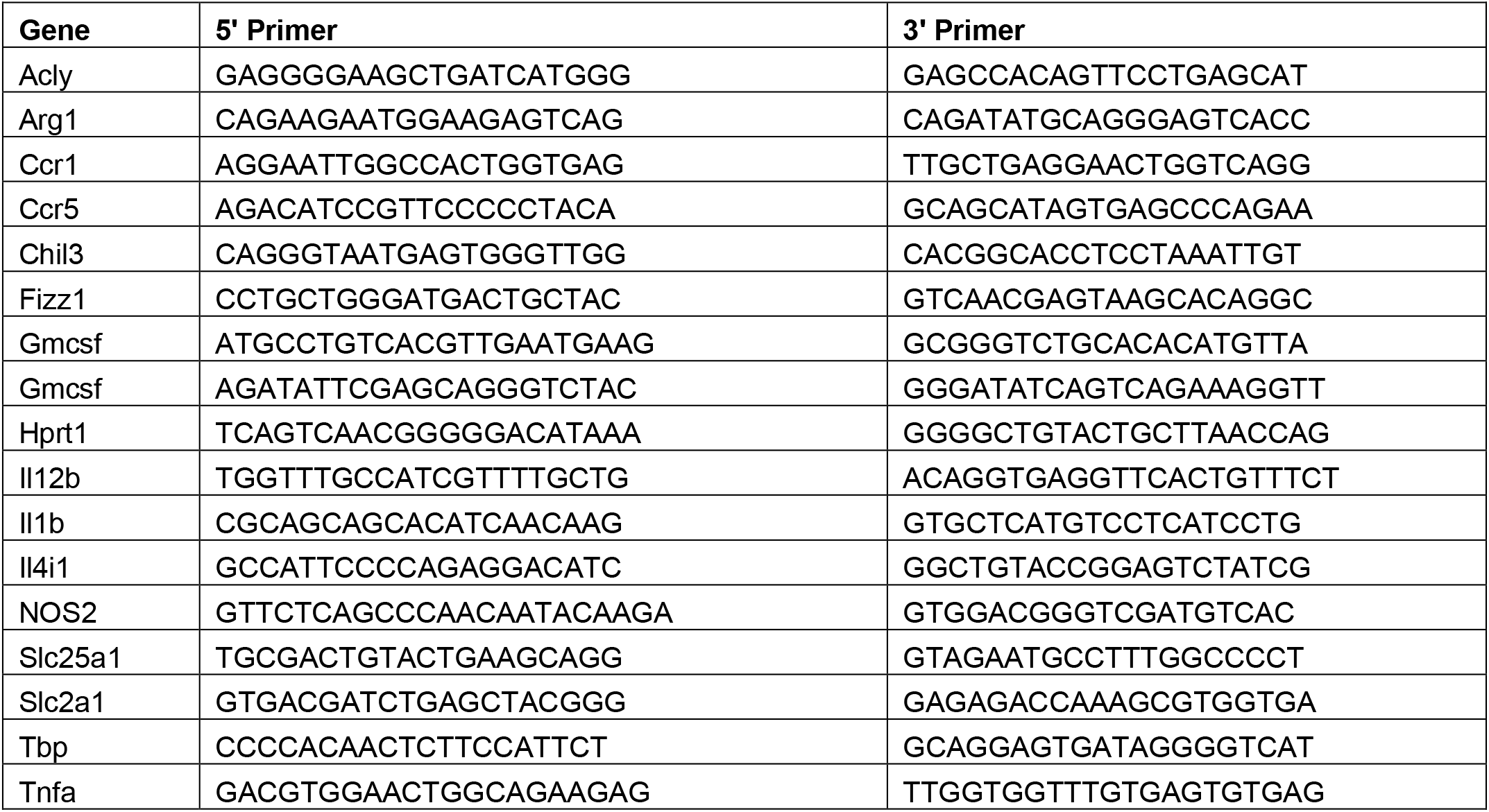

### Proteomics

#### Sample preparation

Six total samples from six macrophage subtypes were prepared in duplicate for mass spectrometry (MS)-based proteomics. The supernatant of each sample’s cell lysate was collected to obtain >70ug of total protein or a protein concentration of 2 ug/ul per sample. Samples were stored at −80 °C until the proteomics experiments.

#### Tandem mass tag (TMT) labeling

Tandem mass tag (TMT) labeling was performed for mass spectrometry (MS) using the TMT 6-plex isobaric labeling kit (ThermoFisher, Cat #90111) according to the manufacturer’s protocol. Two TMT kits were used to label the duplicate sets of the 6 macrophage samples. 60 μg of protein lysates from each sample were reduced with 5 mM dithiothreitol (DTT) at 45 °C for 1 hour, followed by alkylation with 15 mM 2-chloroacetamide at room temperature for 30 minutes. Proteins were precipitated by adding 6 volumes of cold acetone and incubating at −20 °C overnight and were later pelleted by centrifuging at 8000 × g at 4 °C for 10 minutes. Supernatants were discarded, and pellets were resuspended in 100 μl of 100 mM triethylammonium bicarbonate (TEAB) and subsequently digested overnight at 37 °C by adding 1.1 μg of sequencing-grade modified porcine trypsin (Promega, V5113). TMT reagents were reconstituted in 40 μl of anhydrous acetonitrile, and digested peptides were transferred to the TMT reagent vial and incubated at room temperature for 1 hour. The reaction was quenched with 8 μl of 5% hydroxylamine and incubated for 15 minutes. For each of the two TMT experiments, the six samples were combined and dried, followed by 2-dimensional (2D) separation, with the first dimension containing an aliquot from each sample mixture (100 μg). The samples then underwent fractionation using a high pH reverse phase fractionation kit according to the manufacturer’s protocol (Pierce). Fourteen fractions per experiment were dried and reconstituted in 10 μl of loading buffer, 0.1% formic acid and 2% acetonitrile.

#### Liquid chromatography (LC)-MS3 analysis

For raw data acquisition from a total of 28 runs (14 in duplicate), an Orbitrap Fusion (ThermoFisher) and Rapid Separation Liquid Chromatography (RSLC) Systems UltiMate 3000 nano-Ultra Performance Liquid Chromatography (UPLC) (Dionex) were used. To increase accuracy and confidence in protein abundance measurements, a multinotch-MS3 analysis method was employed for MS data analysis. Two microliters from each fraction were resolved in 2D on a nanocapillary reverse phase column (Acclaim PepMap C18, 2 micron particle size, 75 μm diameter × 50 cm length, ThermoFisher) using a 0.1% formic/acetonitrile gradient at 300 nL/min (2–22% acetonitrile in 150 min, 22–32% acetonitrile in 40 min, 20 minute wash at 90% acetonitrile, followed by 50 minute re-equilibration) and sprayed directly onto the Orbitrap Fusion with EasySpray (ThermoFisher; Spray voltage (positive ion) = 1900 V, Spray voltage (negative ion) = 600 V, method duration = 180 minutes, ion source type = nanoelectrospray ionization (NSI)). The mass spectrometer was set to collect the MS1 scan (Orbitrap; 120 K resolution; automatic gain control (AGC) target 2 × 105; max injection time (IT) 100 ms), and then data-dependent Top Speed (3 sec) MS2 scans (collision-induced dissociation; ion trap; NCD 35; AGC 5 × 103; max IT 100 ms). For multinotch-MS3 analysis, the top 10 precursor ions from each MS2 scan were fragmented by high-energy collisional dissociation (HCD), followed by Orbitrap analysis (NCE 55; 60 K resolution; AGC 5 × 104; max IT 120 ms; 100-500 m/z scan range).

#### TMT quantification and data analysis

Raw MS data pre-processing and TMT protein quantification were performed using MSFragger^46^ (peptide identification), the Philosopher toolkit^47^ (peptide validation and protein inference, FDR filtering, and extraction of quantitative information from MS scans) and TMT-Integrator (protein quantification and normalization) as previously described.^48^ A total of 6,919 proteins were quantified and 5,437 proteins were common in the two TMT experiments. The mean and median of Pearson correlation coefficients between the abundance profiles of individual proteins in both TMT datasets were 0.68 and 0.84, respectively. There were 3,631 proteins whose abundance profile correlations were greater than the mean, which we considered consistent between the two TMT experiments. For downstream analysis, the mean-centered normalized data were used. The candidate markers of differentially abundant proteins for each macrophage subtype were identified by a one-tailed t-test for each direction of up- and/or down-regulation against the remaining subtypes with a p-value threshold of 0.001. No multiple testing correction was made in favor of downstream functional analysis.

### Metabolite sample preparation

Intracellular metabolite fractions were prepared from cells grown in non-tissue culture-treated 6-well plates (Corning) that were lysed with cold (−80°C) 80% methanol, then clarified by centrifugation. Metabolite levels of intercellular fractions were normalized to the protein content of a parallel sample, and all samples were dried via speed vac after clarification by centrifugation. Media samples were prepared by collecting 200µl of conditioned or basal media and adding to 800µl of cold 100% methanol. The resultant was clarified by centrifugation and lyophilized via speed vac. Dried metabolite pellets from cells or media were re-suspended in 35 μL 50:50 HPLC grade methanol:water mixture for metabolomics analysis.

### Metabolomics

Agilent 1290 UHPLC and 6490 Triple Quadrupole (QqQ) Mass Spectrometer (LC-MS) were used for label-free targeted metabolomics analysis, as described previously^49^. Agilent MassHunter Optimizer and Workstation Software LC-MS Data Acquisition for 6400 Series Triple Quadrupole B.08.00 was used for standard optimization and data acquisition. Agilent MassHunter Workstation Software Quantitative Analysis Version B.0700 for QqQ was used for initial raw data extraction and analysis. For RPLC, a Waters Acquity UPLC BEH TSS C18 column (2.1 × 100mm, 1.7µm) was used in the positive ionization mode. For HILIC, a Waters Acquity UPLC BEH amide column (2.1 × 100mm, 1.7µm) was used in the negative ionization mode. Further details are found in our previous study^49^.

### Bioinformatics and Statistical Analysis

Bioinformatics analyses were performed using R/Bioconductor. Differential expression or abundance analysis for either up- or down-regulation was done using a one-tailed t-test for each subtype against all the others. A Kolmogorov-Smirnov test did not yield meaningful results across all omics data for downstream analyses. Differential markers were identified using a p-value threshold of 0.001. P-values were not adjusted for multiple testing in favor of flexibility in downstream analyses and biological interpretations. Heat maps were made using R and Morpheus (https://software.broadinstitute.org/morpheus). Metabolomics pathway analyses were performed using MetaboAnalyst 5.0. Bar plots were created using Graph Pad Prism 9. Statistical analyses were performed using Graph Pad Prism 9. Comparisons of two groups were analyzed using unpaired, two-tailed student’s t-test. Comparisons with more than two groups were analyzed with one-way analysis of variance (ANOVA) with Turkey post hoc test. All error bars represent mean with standard deviation.

## Author Contributions

SMB, HL, CJH, and CAL conceived of and designed this study and planned and guided the research. SMB, HL, CJH, and CAL wrote the manuscript. SMB, HL, NGS, LZ, PS, AA, MW, VB, YZ, AIN, MPdM, CJH, and CAL provided key reagents, performed experiments, analyzed, and interpreted data. CAL provided resources, funding, and supervised the research. All authors reviewed and approved the final manuscript.

## Acknowledgements

YZ was supported by R50CA232985. AIN was supported by the NCI U24CA210967. CJH was supported by F32CA228328, K99/R00CA241357, P30DK034933, P30CA062203 and a NPF Research Grant. CAL was supported by the NCI (R37CA237421, R01CA248160, R01CA244931) and UMCCC Core Grant (P30CA046592). Metabolomics studies performed at the University of Michigan were supported by NIH grant DK097153, the Charles Woodson Research Fund, and the UM Pediatric Brain Tumor Initiative.

---

**Supplementary Figure 1.**
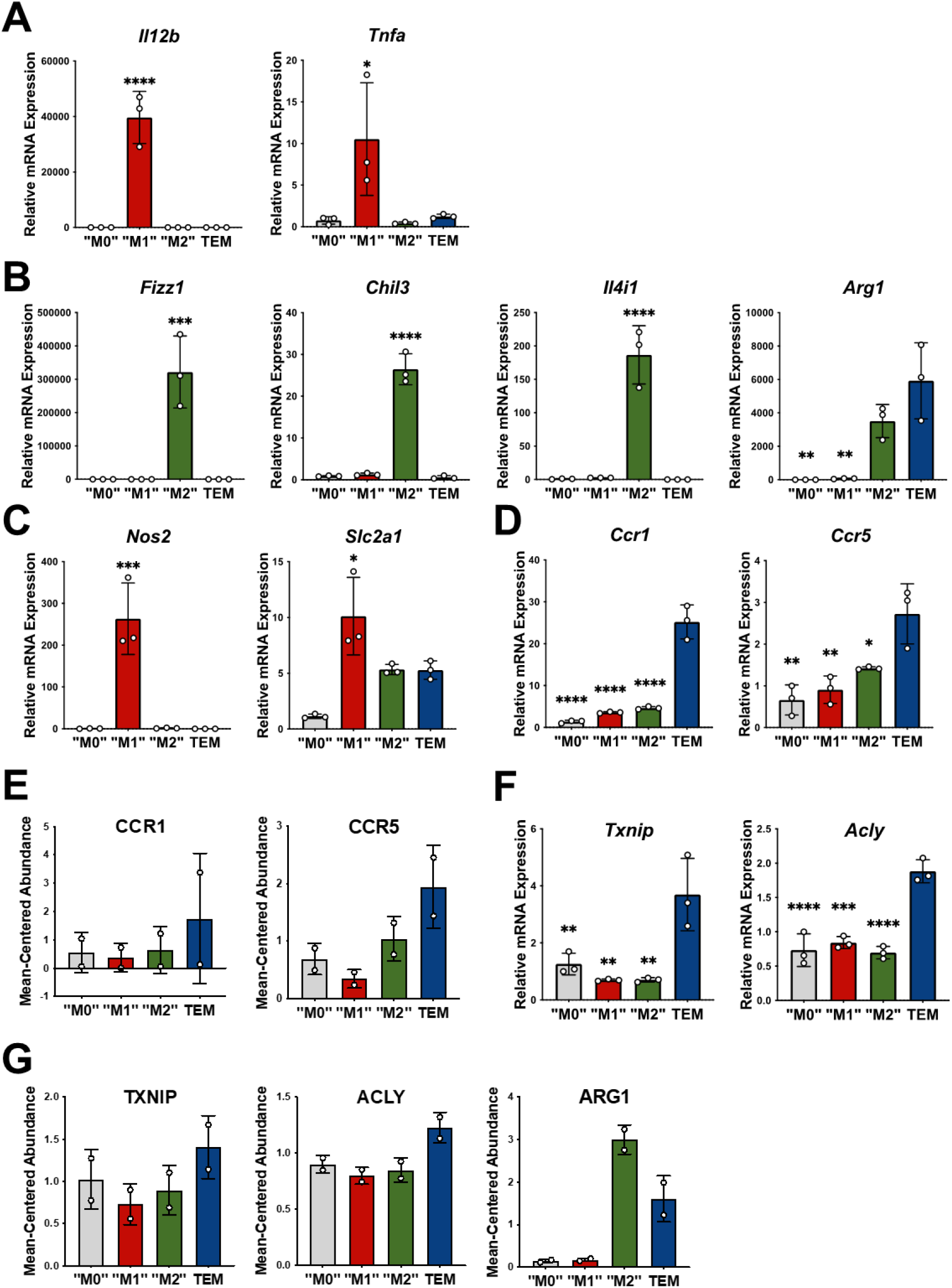
Macrophage marker qPCR validation. **A**) qPCR of classic M1 markers Il12b and Tnfa in M0, M1, M2, and TEM subtypes. ^†^ **B**) qPCR of classic M2 markers Fizz1, Chil3, Arg1, and Il4i1 in M0, M1, M2, and TEM subtypes. ^†^ **C**) qPCR of Nos2 and Slc2a1 in M0, M1, M2, and TEM subtypes. ^†^ **D**) qPCR of TEM cytokine signatures Ccr1 and Ccr5 in M0, M1, M2, and TEM subtypes. ^†^ **E**) Proteomics mean-centered abundance of CCR1 and CCR5 in M0, M1, M2, and TEM subtypes. **F**) qPCR of TEM enzyme markers Txnip and Acly in M0, M1, M2, and TEM subtypes. ^†^ **G**) Proteomics mean-centered abundance of TXNIP, ACLY, and ARG1 in M0, M1, M2, and TEM subtypes. ^†^Significance comparisons are relative to TEM subtype (* P < 0.05; ** P < 0.01, *** P < 0.001, **** P < 0.0001).

**Supplementary Figure 2.**
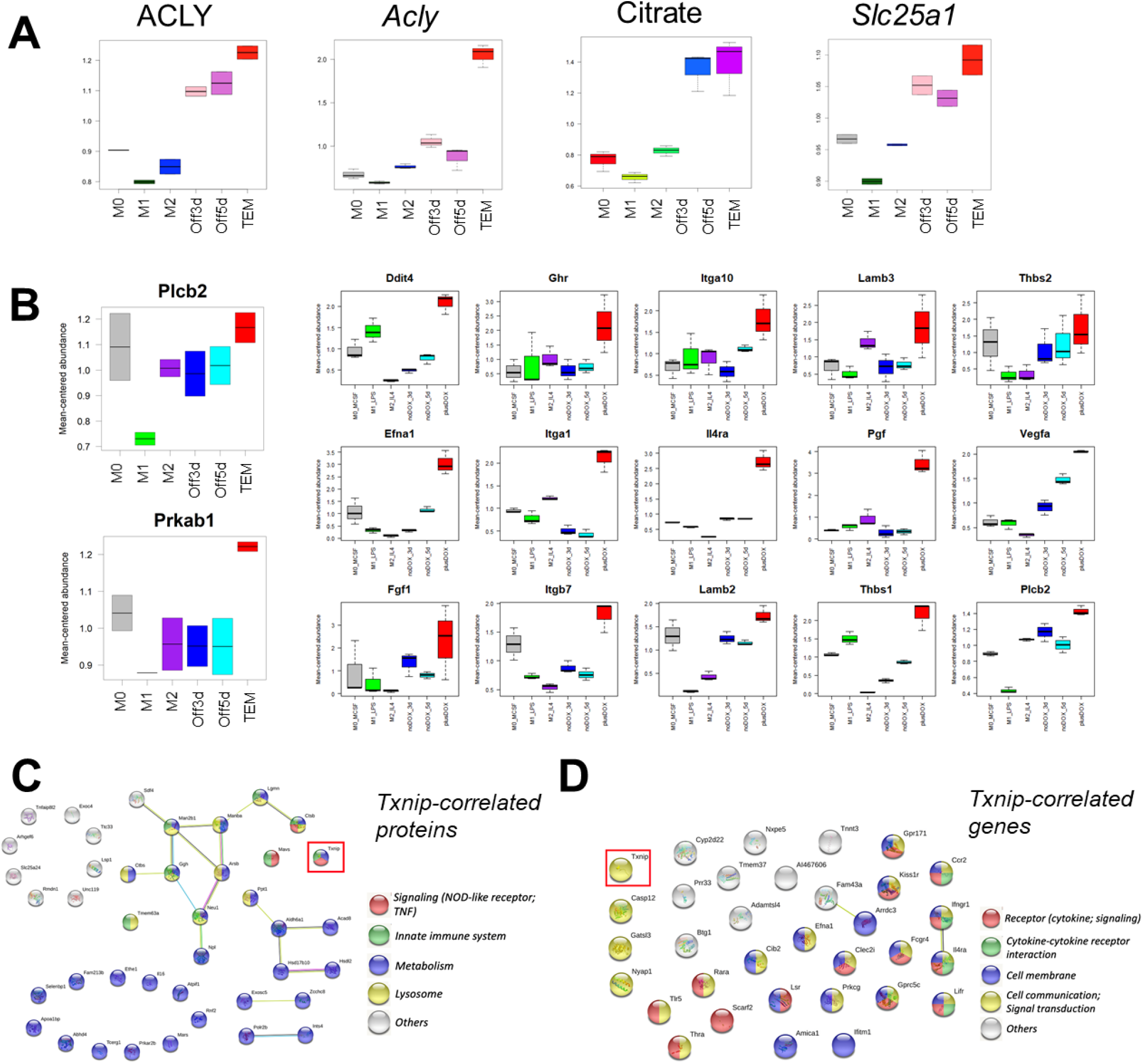
Multiomic pathway analyses. **A**) Proteomics and transcriptomics box plot of Acly, metabolomics box plot of citrate, and transcriptomics box plot of Slc25a1 for M0, M1, M2, Kras-Off 3-day, Kras-Off 5-day, and Kras-On TEM macrophage subtypes.* **B**) Proteomics box plots of PLCB2 and PRKAB1 and transcriptomics box plots of 15 PI3K-related genes.* **C**) STRING-based pathway analysis of Txnip-correlated proteins in TEM. **D**) STRING-based pathway analysis of Txnip-correlated genes in TEM. * “Off3/5d” = “noDOX_3/5d” = Kras-Off TEM subtype; “plusDOX” = Kras-On TEM subtype.

**Supplementary Figure 3.**
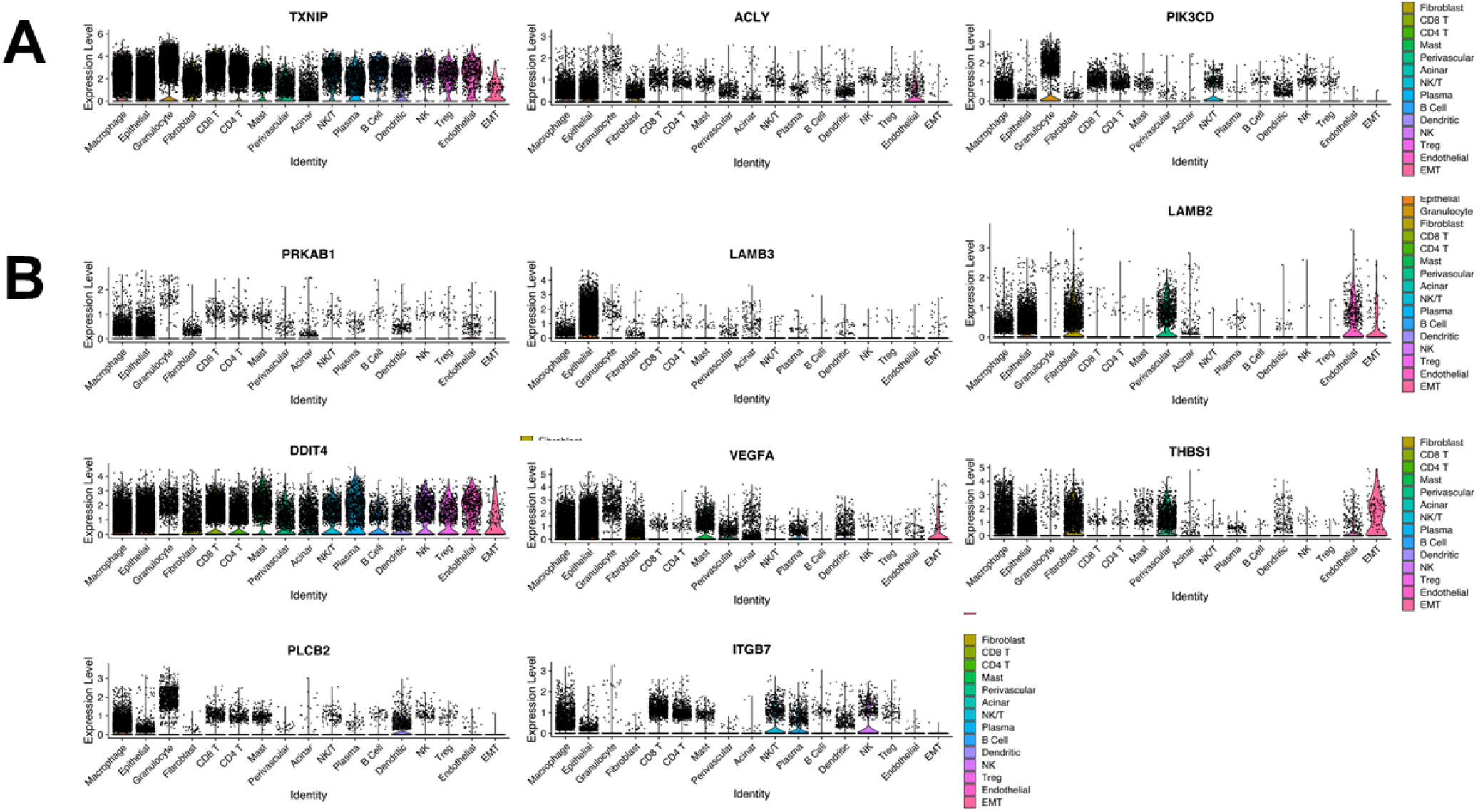
Expression of pancreatic TEM markers in human tumors. **A**) Violin plots of TEM markers Txnip, Acly, and Pik3cd expressed in human pancreatic tumors. **B**) Violin plots of PI3K-related genes expressed in human pancreatic tumors (Prkab1, Lamb3, Lamb2, Ddit4, Vegfa, Thbs1, Plcb2, Itgb7).

**Supplementary Figure 4.**
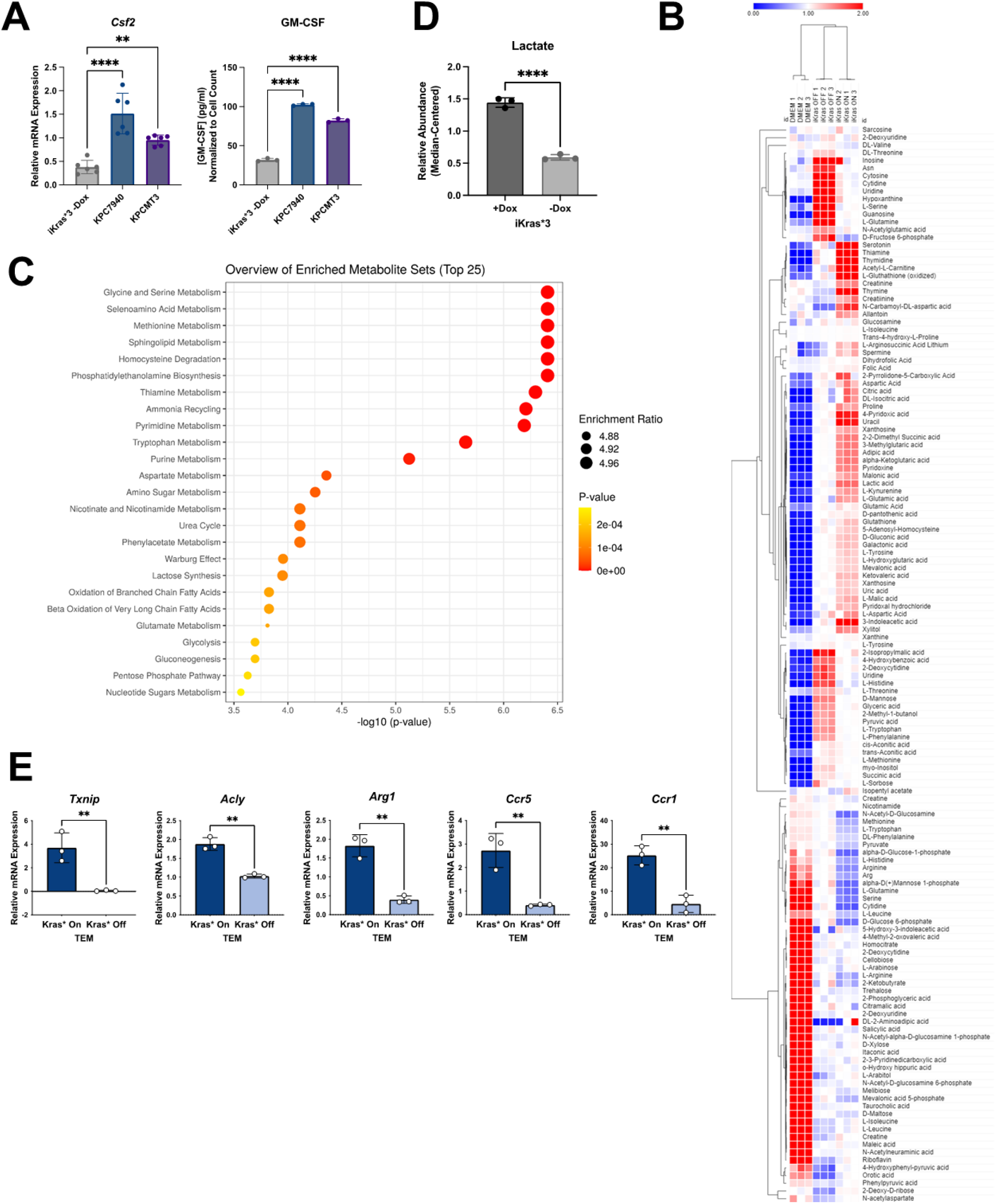
Effects of mutant Kras on the extracellular environment. **A**) qPCR of Csf2 expression and ELISA of GM-CSF release in Kras-extinguished iKras*3 cells and two KPC PDA lines, KPCMT3 and KPC7940. **B**) Heat map of significant (P < 0.01) extracellular metabolites released from Kras-expressing vs. Kras-extinguished iKras*3 cells. **C**) Enrichment analysis of Kras-expressing vs. Kras-extinguished iKras*3 extracellular metabolomics. **D**) Extracellular lactate abundance from iKras*3 cells +/-dox as measured by enzymatic fluorescence assay. **E**) qPCR validation of TEM enzyme and cytokine markers Txnip, Acly, Arg1, Ccr5, and Ccr1 in Kras-On vs Kras-Off TEMs. (* P < 0.05; ** P < 0.01, *** P < 0.001, **** P < 0.0001).

**Supplementary Figure 5.**
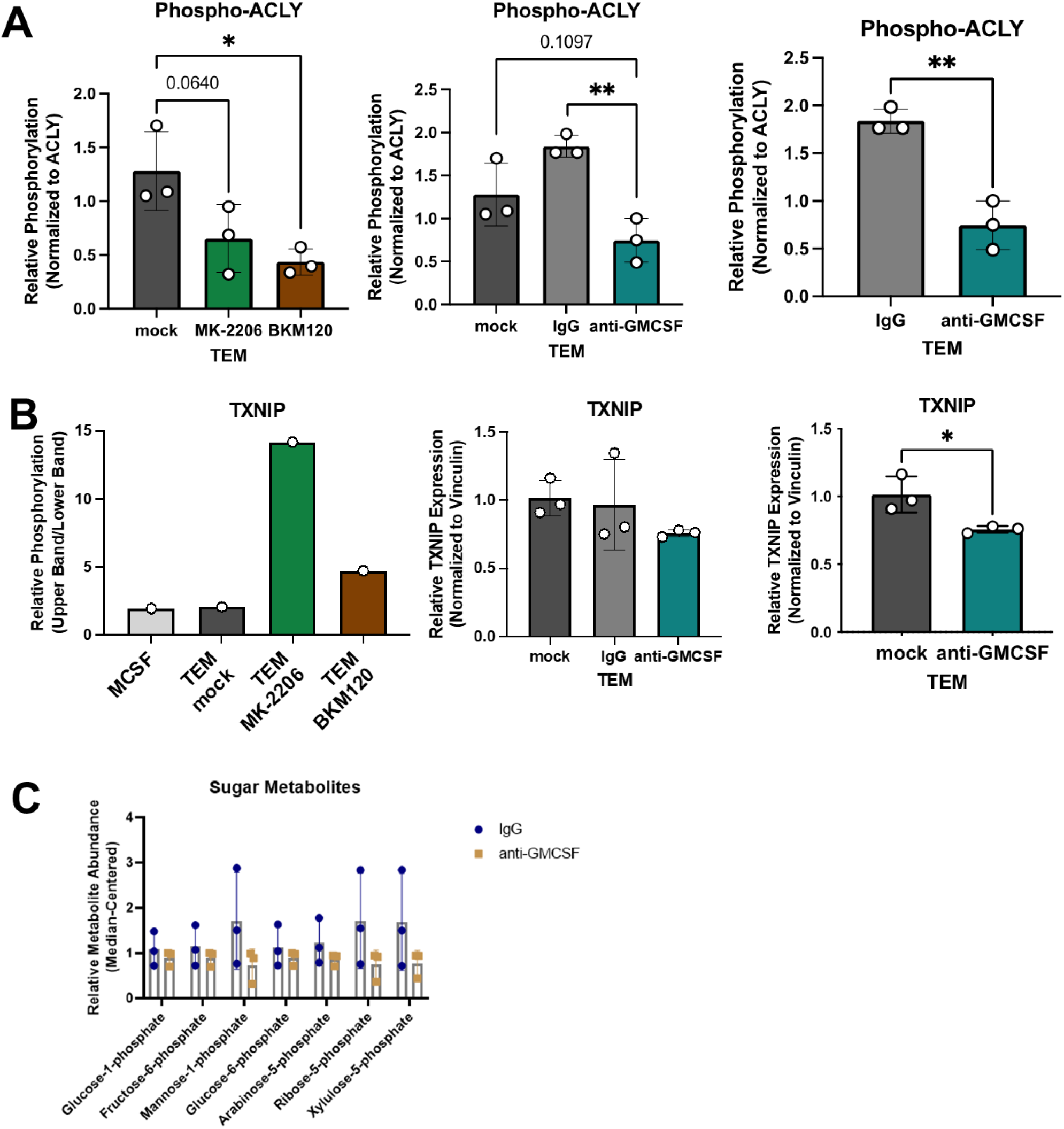
GM-CSF neutralization disrupts the TEM metabolic phenotype. **A**) Bar graphs of pACLY quantitation relative to ACLY expression in TEM mock, MK-2206, BKM120, IgG, and anti-GMCSF groups*. **B**) Bar graphs of TXNIP phosphorylation quantitation from Fig. 5A and TXNIP expression quantitation in TEM mock, IgG, and anti-GMCSF groups*. **C**) Bar graph of phosphorylated sugar metabolites in IgG and anti-GMCSF TEM groups. (* P < 0.05; ** P < 0.01). *Replicates represent western blots performed on independent repeat experiments.

